# Multimodal cortical connectome in the default mode network across the adult lifespan

**DOI:** 10.1101/2025.05.16.654309

**Authors:** San-wang Wang, Xiaobo Liu, Jinzhi He, Xiang-wen Chang, Qiu-xuan Yu, Bin Wan, Zhen-Qi Liu, Wei-feng Mi, Le Shi

**Author notes:** Corresponding Author: Le Shi, PhD, Institute of Mental Health/Peking University Sixth Hospital, 51 Huayuanbei Road, Haidian District, Beijing, 100191, China, Wei-feng Mi, MD, Institute of Mental Health/Peking University Sixth Hospital, 51 Huayuanbei Road, Haidian District, Beijing, 100191, China. These authors contributed equally to this work (San-wang Wang, Xiaobo Liu).

## Abstract

The default mode network (DMN) critically underpins cognitive and affective functions throughout the adult lifespan; however, detailed insights into its complex neuroarchitecture and connectivity patterns across aging remain limited. Leveraging the open-access CamCAN dataset, comprising structural and diffusion magnetic resonance imaging (MRI) alongside magnetoencephalography (MEG) data from 599 adults spanning ages 18 to 88 years, we systematically investigated age-associated changes in multimodal connectomes within the DMN. Our analyses revealed a progressive decline in both structural and functional connectivity among DMN subregions with advancing age. Additionally, MEG-based connectivity assessment demonstrated age-related decreases in high-frequency oscillatory activity (alpha, beta, gamma bands) accompanied by increases in low-frequency oscillations (theta band). Integrating structural data with neurophysiological measures further revealed age-dependent shifts in neurophysiological-structural coupling within the prefrontal cortex, characterized by strengthened coupling at theta frequencies but weakened coupling at higher frequencies. Conversely, coupling within the posterior cingulate cortex consistently declined across all examined frequency bands. Notably, theta-band coupling within the prefrontal cortex significantly correlated with age-related memory performance variations. Collectively, our findings delineate nuanced changes in DMN information transmission dynamics across adulthood, underscoring its promise as a neurobiological biomarker reflective of cognitive aging heterogeneity.

## Introduction

The default mode network (DMN) is a highly active neural network during rest, involving key regions such as the prefrontal cortex (PFC), posterior cingulate cortex (PCC), temporal regions, lateral parietal cortex, and hippocampus^1,2^. Despite being widely distributed across the cortex, these DMN regions are interconnected, supporting abstract cognitive functions and underpinning core mechanisms of consciousness^3^. While cognitive functions are known to evolve across the lifespan, the manner in which the cortical connectome within the DMN changes with age remains largely unexplored.

The DMN, with its extensive cortical distribution, is essential for integrating basic information and supporting higher-order cognitive functions. However, older adults show weakened structural and functional connectivity within the DMN, particularly between the medial prefrontal cortex and the posterior cingulate cortex, during cognitive tasks such as working memory, episodic encoding, and semantic classification^4,9-11^. These findings underscore the reduced ability of older adults to allocate cognitive resources effectively. Furthermore, in older adults experiencing subjective cognitive decline, disrupted functional connectivity between the posterior DMN and the para-hippocampal gyrus is linked to a higher risk of Alzheimer’s disease (AD) ^12^. Longitudinal studies have revealed non-linear age-related changes in DMN functional connectivity, with early increases potentially serving as biomarkers for cognitive aging^13^. Together, these findings underscore the importance of examining the DMN and its internal connectivity to uncover the mechanisms behind the dynamic changes in brain network functioning across the adult lifespan.Thus, we hypothesize that the DMN would serve as a crucial framework for comprehending the intricate neurodynamic changes that occur during the adult lifespan.

Most studies of previous studies predominantly utilized single-modality imaging to reveal changes in the DMN’s internal brain the functional connectome (FC) or a structural connectome (SC) cross the lifespan. Currently, the internal connections between brain structure and function are receiving increasing attention and research, as the anatomy and physiology of the brain are inextricably linked^14^. SC-FC coupling of DMN has been found to correlate with cognitive integrity significantly but decreases with age^15^. The structural connectivity is calculated based on the white matter integrity, and traditional FC is calculated based on functional, that relies on blood oxygen level-dependent (BOLD) signals, which indirectly measure ultra-low frequency brain activity, neglecting high-frequency signals and the structural basis of these transmissions^16^. However, high-frequency electrophysiological activities and brain structure are crucial for complex cognitive functions^17^. Recent EEG and MEG studies have shown age-related physiology changes within the DMN regions, respectively, between ^18-21^. Therefore, it is important to integrate structural and electrophysiological modalities to systematically clarify the effects of aging on the DMN from electrophysiological and structural perspectives. Coupling electrophysiological measurements with structural MRI is considered a new metric for measuring brain communication efficiency linked to various cognitive abilities^22,23^. For instance, previous studies have shown that structure-function coupling is related to age-related cognitive decline^24^. A multimodal study focusing on the DMN’s internal connectivity patterns is urgently needed to systematically elucidate the mechanisms of age-related changes across the adult lifespan.

In this study, we comprehensively analyzed the relationship between DMN, age, and memory using multimodal data from the Cambridge Centre for Ageing and Neuroscience (Cam-CAN) database, comprising 600 individuals aged between 18 and 88. ^25^. Utilizing MEG and DTI, we first constructed neurophysiological connectome (NC) and SC within the DMN. Through coupling analysis, we revealed the joint patterns of SC-NC across the lifespan. Furthermore, we elucidated the relationship between structural integrity and brain dynamics in the context of aging and memory performance, highlighting the significance of SC-NC coupling as a potential biomarker for cognitive changes associated with age.

## Method

### Cambridge Centre for Aging and Neuroscience Project Dataset

In this study, participants were 608 healthy adults (307 males/301 females, the average age was 54.19 ± 18.19), including resting state MEG and MRI from Cambridge Centre for Ageing and Neuroscience Project Dataset^25^. The resting state MEG were acquired using a 1000 Hz sampling rate MEG system. Also, to test the cognition relevance of kinetic properties, we used the visual short-term memory capacity from the CAM-CAN dataset, consisting included in this study.

### fMRI data preprocessing and connectome construction

The fMRI data underwent conversion from raw DICOM files to the Brain Imaging Data Structure (BIDS) format using HeuDiConv v0.13.1. Subsequent structural and functional preprocessing was performed with fMRIPrep 23.0.2^1^, leveraging Nipype 1.8. as the processing pipeline foundation^2^. Anatomical preprocessing included a series of steps: intensity normalization, brain extraction, tissue segmentation, surface reconstruction, and spatial normalization. Functional preprocessing encompassed head motion correction, slice-time correction, and co-registration. The functional time series were parcellated using the Schaefer 200×7 atlas^3^ and underwent confound removal following the “simple” strategy^4^. This strategy incorporated high-pass filtering, motion and tissue signal removal, detrending, and z-scoring. Functional connectivity matrices were subsequently computed for each participant by calculating zero-lag Pearson correlation coefficients.

### Structural data preprocessing and connectome construction

Diffusion MRI preprocessing was performed using QSIprep 0.17.0^33^. The diffusion-weighted images underwent denoising, Gibbs unringing, bias field correction, intensity normalization, head-motion estimation and Eddy current correction, registration, and normalization to standard space. Structural connectivity matrices were reconstructed using the “mrtrix_multishell_msmt_ACT-hsvs” pipeline. Fiber orientation distributions were estimated using the MSMT-CSD algorithm. Probability tractography was conducted using the iFOD2 probabilistic tracking method with anatomically constrained tractography (ACT) based on T1-weighted segmentation constraints. Streamline weights were calculated using the SIFT2 algorithm to construct the structural connectivity matrix (Figure 1c).

**Figure 1.**
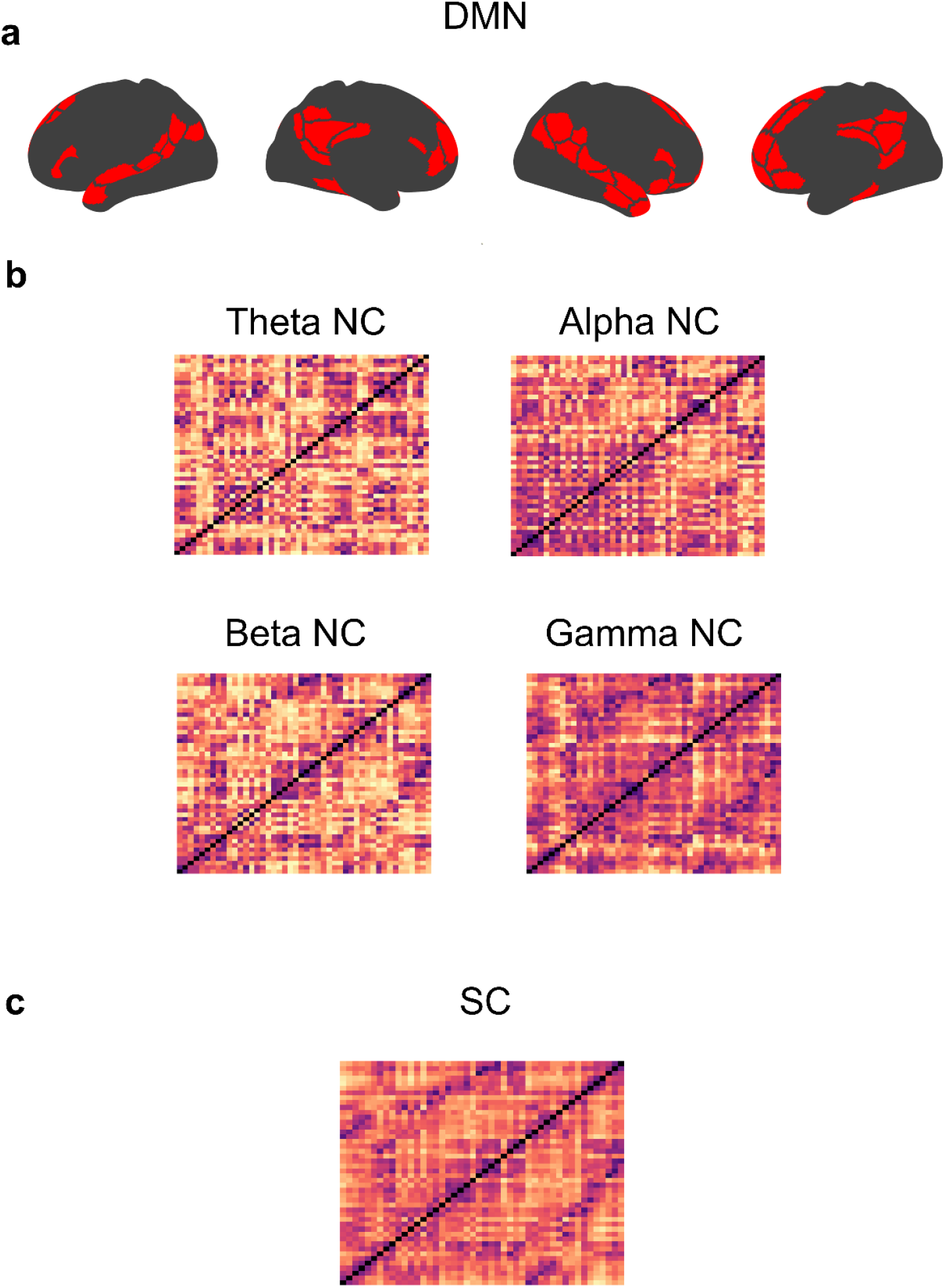
Multimodal connectome among sub-regions in DMN. a.Sub-regions of DMN using Schaefer 200 parcellation. b. Group-average MEG neurophysiological connectome within DMN across 4 frequency bands, theta (4-8 Hz), alpha (8-13 Hz), beta (13-30 Hz), gamma(30-50Hz). c. Group-average structural connectome among subregions in DMN.

### MEG data preprocessing and connectome construction

All data analysis was conducted via Brainstorm with default parameters unless specified^26^ and adhered to the good-practice guidelines^27^. The slow wave and DC-offset components were initially removed from the raw signal using a high-pass FIR filter with a 0.3-Hz cutoff. Raw MEG signals were visually inspected to eliminate bad channels and data segments. Line noise artifacts (60 Hz) and their harmonics were filtered out using a notch filter. Subsequently, a low-pass filter (0.6 – 280 Hz) was applied to eliminate high-frequency noise while retaining most of the signal. Signal-space projection (SSP) techniques were employed to identify and remove heartbeats and eye-blinking artifacts based on electro-cardiogram and electrooculogram recordings.

The clean data were segmented into 30-second epochs^28^ to ensure robust resting-state results. These segments were then filtered to the desired frequency range and resampled at four times the cutoff frequency to preserve most of the data and prevent aliasing^29^.

Clean MEG sensor data were utilized for source localization. Individual T1-weighted MRIs were acquired for co-registration with MEG data (1.5T Siemens Sonata). Tissue segmentation and cortical surface extraction were performed using FreeSurfer (http://surfer.nmr.mgh.harvard.edu/)^30^. Cortical surfaces were tessellated into triangular meshes, and inflated surfaces were used to visualize cortical data. Co-registration with MEG sensor locations was achieved using digitized head points collected during the MEG session. The forward head models were obtained using the overlapping spheres approach, and the cortical source model was computed using linearly constrained minimum-variance (LCMV) beamforming, all within Brainstorm (2016 version). Noise covariance matrices, estimated from empty-room MEG recordings, were applied to normalize estimated source variance and mitigate source depth effects.

The DMN cortical signals were parcellated (Figure 1a) using the Schaefer 200 template via principal component analysis ^31^. To calculate neurophysiological connectome in various frequency band, we implemented time frequency analysis, which were calculated using amplitude envelope correlation (ACE) with orthogonalized signals to address signal leakage for each frequency band^532^. Then further analyses were performed across five frequency bands, including Theta (4 – 8 Hz), Alpha (8 – 13 Hz), Beta (13 – 30 Hz), and Gamma (30 – 50 Hz), via averaging the connectome matrix within the same frequency bands (Figure 1b).

### Graph theory analysis

To explore the importance as well as efficiency of sub-regions information exchange within DMN, we implemented graph theoretic indices on connectome patterns via brain connectivity toolbox (BCT, https://sites.google.com/site/bctnet/). Specifically, we calculated node strength and node-local efficiency of DMN subregions for each subject. Note that global efficiency is a network integration measure that describes the overall integration of information flow within a network^34^. It is defined as the average of the inverse shortest path lengths between all nodes in the network.

### Structural-physiological coupling

To explore the relationship between SC and NC, we calculated Pearson’s correlation between SC and FC as well as NC with 4 frequency bands within subregions in DMN.

### Aging association

To explore the impact of aging on DMN, we calculated Pearson’s correlation between FC, SC, and various NC as well as their graph-theoretic indices with age, respectively, sex, and handedness as covariates. Furthermore, we investigated the overlap of consistent network changing patterns across all modalities.

### Visual short-term memory capacity prediction

To elucidate the cognition relevance and various-modality networks, we employed a multivariate modeling approach to predict visual short-term memory capacity via those identical connectomes within DMN. We applied a 10-fold cross-validation approach on all age-related modality connectome to train a support vector regression (SVR) model aimed at predicting memory scores. During each fold, the SVR model (using the LIBSVM library, https://www.csie.ntu.edu.tw/~cjlin/libsvm/) was trained with default settings and a linear kernel^6^. Following training, the predicted scores from each fold were compiled, and prediction accuracy was determined by calculating Pearson’s correlation coefficient between the predicted and actual memory scores. To assess model robustness, the cross-validation procedure was repeated 1,000 times, with the average predicted scores across iterations serving as the result.

## Results

### Functional connectome changes of the DMN across human aging

We first assessed functional connectivity (FC) within the DMN using Pearson correlations derived from fMRI data to identify age-related connectivity changes (Figure 2a). We found that aging was associated with an overall decline in functional connectivity within the DMN (*p*_*FDR*_ < 0.05). Notably, connectivity increased within temporal and prefrontal cortex (PFC) subregions, as well as between temporal and posterior cingulate cortex (PCC) subregions, particularly in the right hemisphere. Conversely, connectivity between PFC and PCC subregions and intra-regional connectivity within these areas decreased significantly with advancing age. This suggests a complex pattern of both strengthening and weakening connections, possibly reflecting compensatory neural adaptations or reductions in integrative processing. Graph theory analyses (Figures 3c and 3d) supported this interpretation, showing decreased node strength but increased node efficiency in PFC and PCC subregions, indicative of potential compensatory mechanisms.

**Figure 2.**
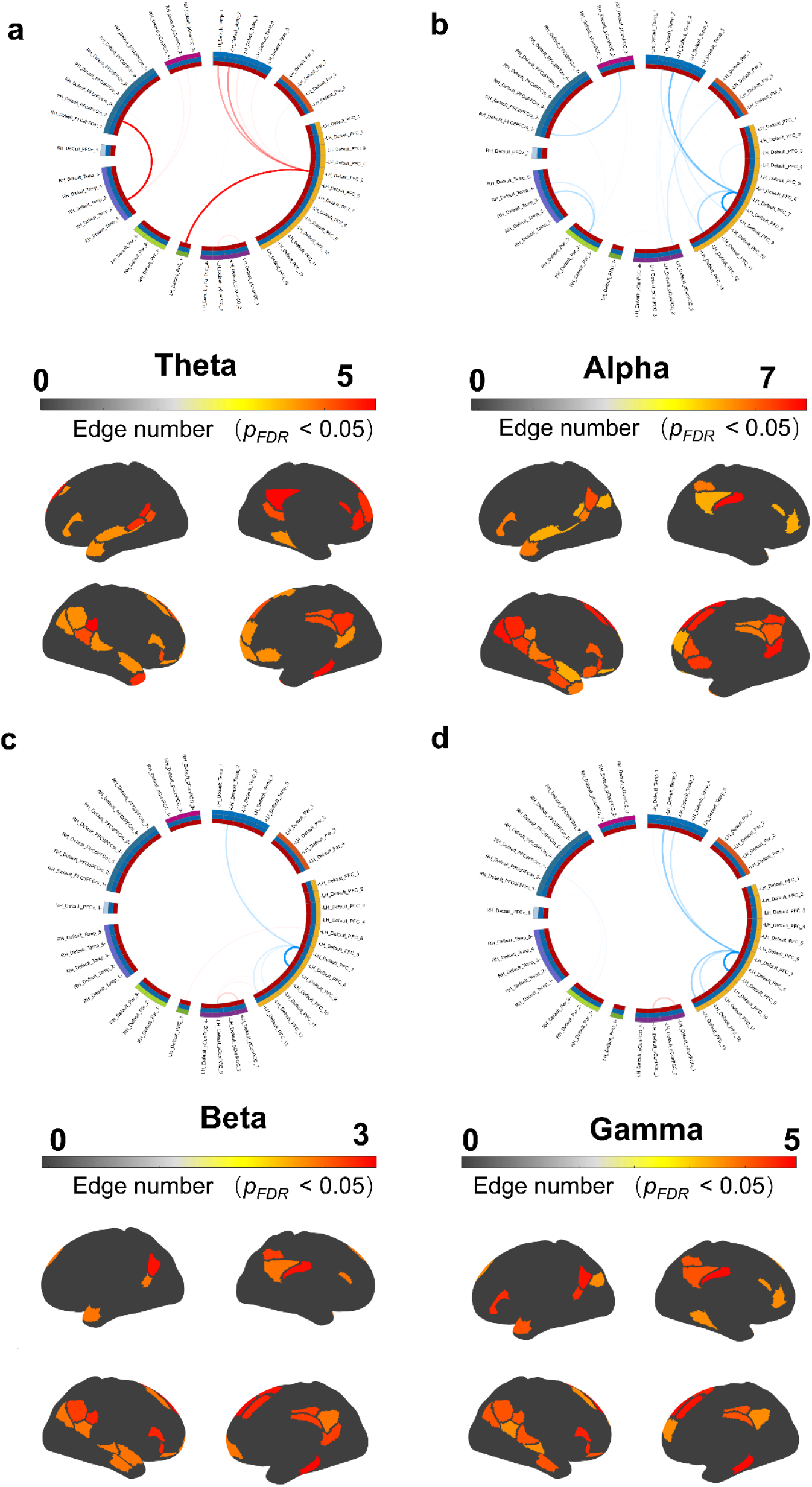
Age-associated neurophysiological DMN networks in 4 frequency bands. Note that red lines represent the edges positively related to age and blue lines represent the edges negatively related to age. A cortical map represents the number of significant edges on the cortex. a. Age related network pattern and their cortical distribution in theta (*p*_*FDR*_ < 0.05). b. Age related network pattern and their cortical distribution in alpha (*p*_*FDR*_ < 0.05). c. Age related network pattern and their cortical distribution in beta (*p*_*FDR*_ < 0.05). d. Age related network pattern and their cortical distribution in gamma (*p*_*FDR*_ < 0.05).

**Figure 3.**
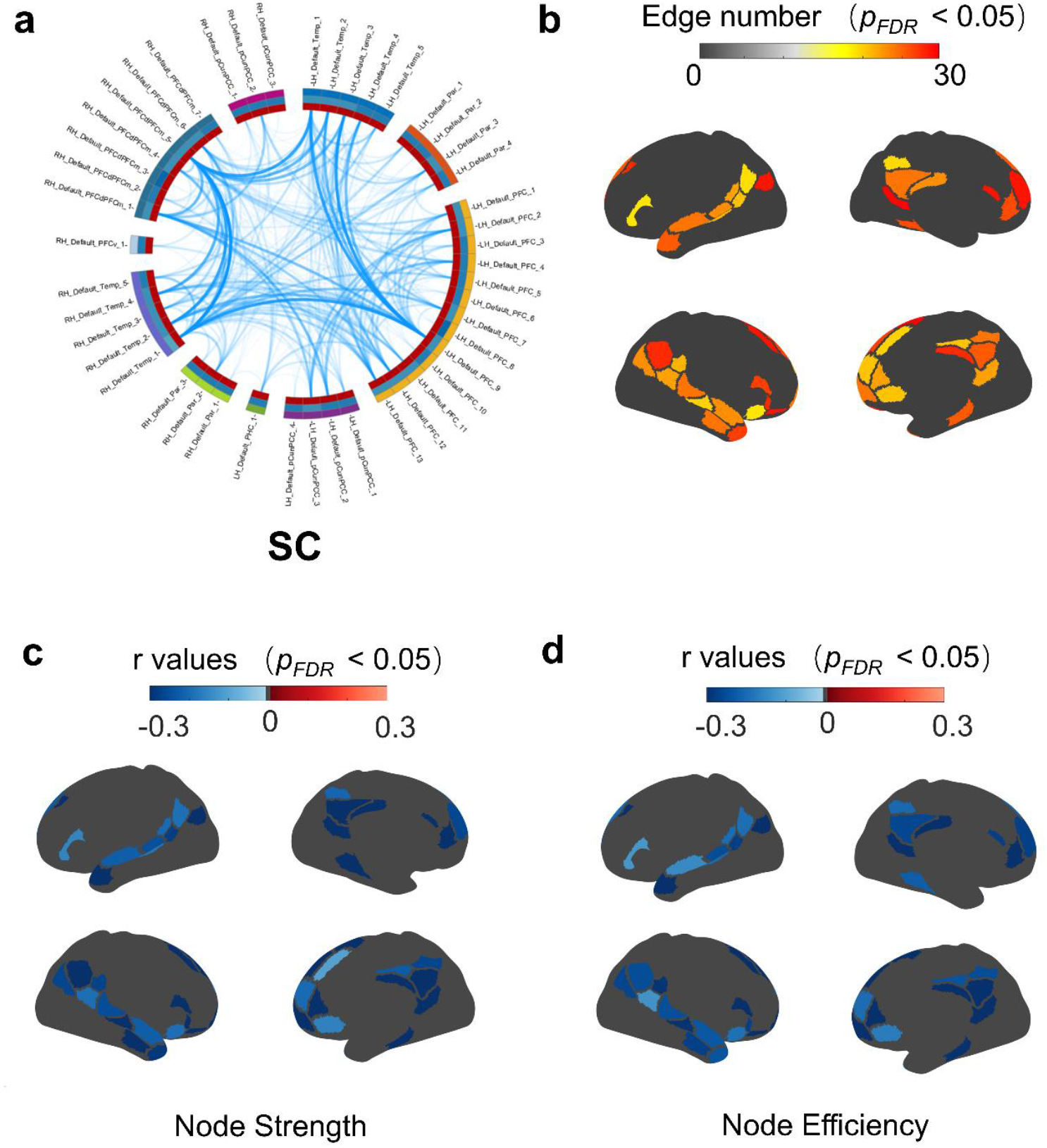
Age-associated structural connectome among subregions in DMN networks. Note that red lines represented the edges positively related to age and blue lines represented the edges negatively related to age. a. Age related network patterns and significant aging regions. b. Correlation between age and node strength within DMN (*p*_*FDR*_ < 0.05). c. Correlation between age and node strength within DMN (*p*_*FDR*_ < 0.05). d. Correlation between age and node efficiency within DMN (*p*_*FDR*_ < 0.05).

### Structural connectome across human aging

We next analyzed structural connectivity (SC) using diffusion MRI and probabilistic tractography to investigate white matter integrity across DMN subregions (Figure 3a, b). Aging significantly reduces structural connectivity in all DMN subregions, most prominently affecting the PFC and PCC (*p*_*FDR*_ < 0.05). Graph theoretical measures further confirmed these findings, showing a consistent decline in node strength and efficiency across the DMN with age (Figures 3c, 3d). These structural declines likely underpin the observed functional connectivity changes, highlighting a critical link between structural integrity and functional network dynamics.

### Neurophysiological connectome changes of DMN with human aging

We first noticed the group-averaged multi-band neurophysiological phrase covariation matrix within DMN, including theta, alpha, beta, and gamma bands. In the theta band (Figure 2a), aging is associated with consistent increased connectivity within the DMN, notably between the PFC subregions and temporal subregions in both hemispheres. In the left hemisphere, connectivity also increased between the PFC subregions and parietal, parahippocampal subcortices, and within PCC. In the right hemisphere, connectivity also increased between the PCC and temporal subregions. In the alpha band (Figure 2b), aging leads to a consistent decrease in connectivity within DMN. Specifically, in the left hemisphere, connectivity within PFC, between PFC subregions and temporal subcortices, between PCC subregions and temporal subcortices, and parietal cortices declined. In the right hemisphere, connectivity between temporal subcortices and parietal subcortex, between PFC and PCC subregions decreased. In the beta band (Figure 2c), connectivity changes are mixed in the left hemisphere, with decreases observed in the prefrontal-temporal connections and within prefrontal connections but increases in connections involving the parahippocampal and prefrontal cortices. In the gamma band (Figure 2d), a significant decrease in connectivity is noted, particularly within the prefrontal and temporal subregions in the left hemisphere, while some increases are seen within PCC subregions. These results suggest that aging induces a neurophysiological tendency for a main decline in high frequencies but increases in theta frequency in DMN connectivity, especially involving left PFC, reflecting the complex effects of aging on brain network dynamics.

### NC-SC coupling within DMN across human aging

We further explored coupling between structural and neurophysiological connectomes (NC-SC coupling) to assess how anatomical changes influence neurophysiological dynamics (*p*_*FDR*_ < 0.05). We observed increased theta-band coupling with advancing age within bilateral PFC, medial temporal cortex, inferior frontal gyrus, and angular gyrus. Higher-frequency coupling (alpha, beta, gamma) showed mixed results, indicating differential adaptations across frequency bands. This frequency-dependent coupling change emphasizes the dynamic interaction between structural deterioration and neurophysiological adaptation during aging.

### Cognition relevance of various connectome within DMN

Figure depiction of multimodal (electrophysiological, functional, and structural) connectivity within the DMN and its predictive capacity for visual short-term memory performance. Figure 5a shows the chord diagram illustrates multimodal connectivity across various subregions of the DMN. Each node represents distinct DMN subregions, including temporal (Temp1 to Temp5), parietal (Par1 to Par4), and PFC1 areas. Node size is proportional to node strength, with larger nodes indicating higher connectivity strength. Connections between nodes reflect statistically significant multimodal associations. Prominent multimodal connectivity patterns are primarily observed between temporal and parietal regions, underscoring the intricate structural and neurophysiological coupling patterns within DMN subregions.

**Figure 4.**
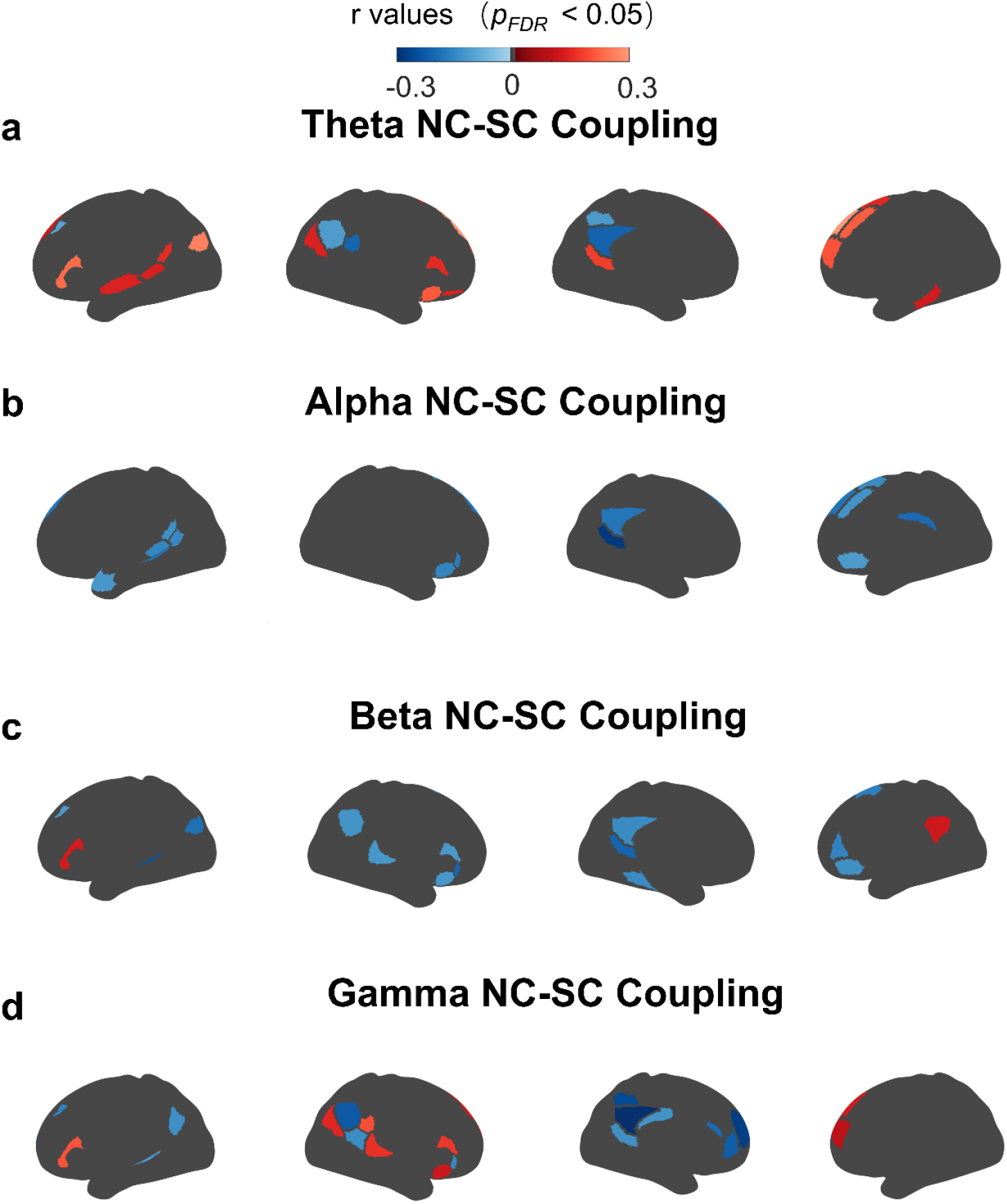
The association between NC-SC coupling and age of DMN networks in 4 frequency bands. a. Age related network pattern of DMN networks in theta frequency band (*p*_*FDR*_ < 0.05). b. Age related network pattern of DMN networks in alpha frequency band (*p*_*FDR*_ < 0.05). c. Age related network pattern of DMN networks in beta frequency band (*p*_*FDR*_ < 0.05). d. Age related network pattern of DMN networks in gama frequency band (*p*_*FDR*_ < 0.05).

**Figure 5.**
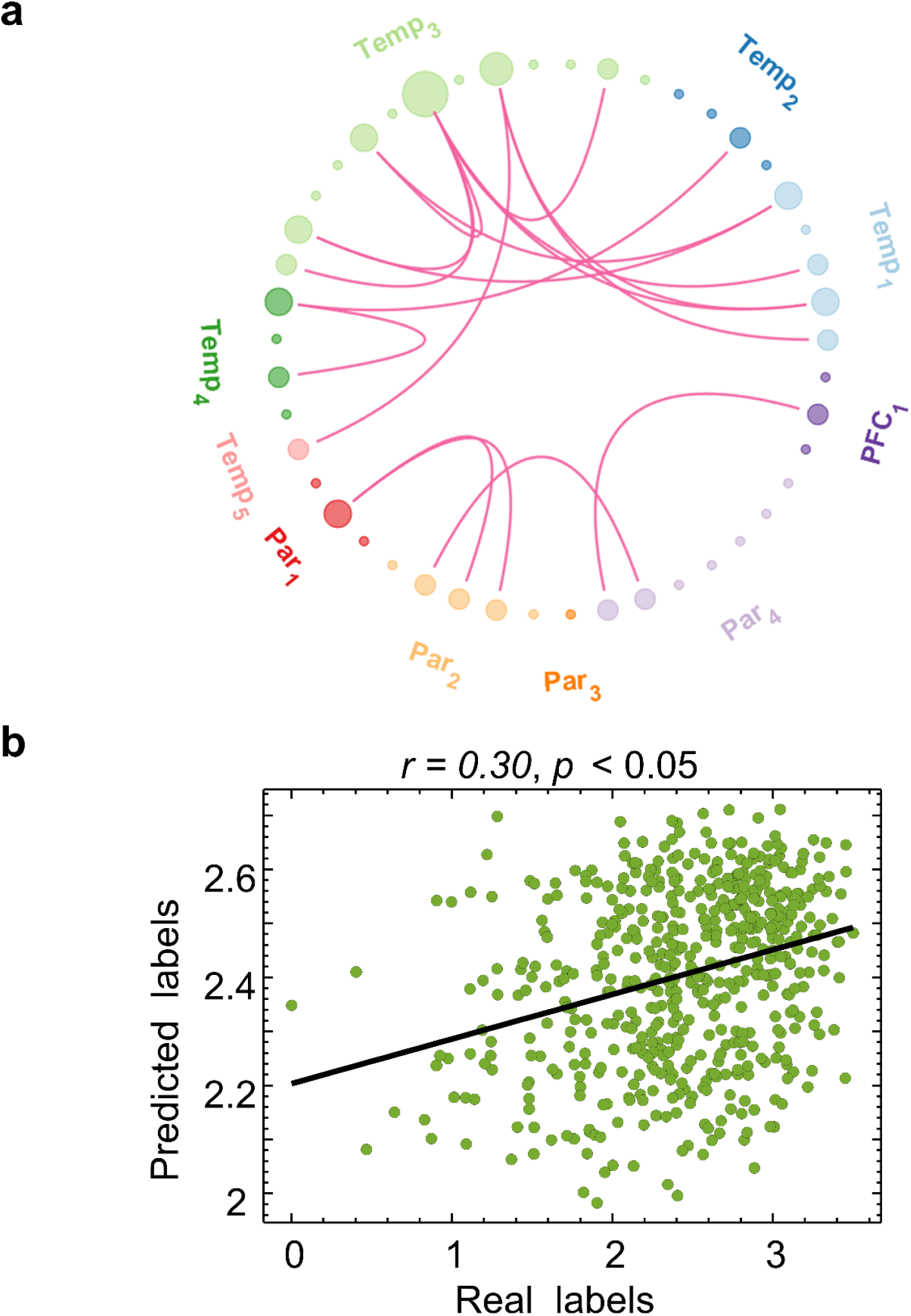
The overlap age-related network predicts visual short-term memory capacity. a. The overlap in age-related networks across all modalities (functional connectome, neurophysiological connectome and structural connectome). b. The multiple modalities overlap network predicted human visual short-term memory capacity (*r* = 0.30, *p* < 0.05).

Figure 5b reveals the scatter plot depicts the prediction of visual short-term memory performance, comparing actual (real) and predicted memory scores. The vertical axis shows predicted visual short-term memory performance scores generated by the model, while the horizontal axis displays the actual visual short-term memory scores. The regression analysis indicates a significant correlation (*r* = 0.30, *p* < 0.05), suggesting that multimodal connectivity patterns within the DMN can significantly predict individual differences in visual short-term memory capacity. Overall, these findings emphasize the significance of multimodal connectivity within the DMN as a potential biomarker for cognitive function, particularly in the context of age-related cognitive decline.

## Discussion

In the present study, we implemented modalities data from a large sample open dataset to investigate the paradigm of the connectome within the default mode network (DMN) across the adult human lifespan. Our findings revealed that in the aging part of the adult lifespan, low-frequency (theta band) connectivity within DMN subregions (PFC, temporal regions, PCC) was enhanced, and high-frequency (alpha, beta, gamma bands) connectivity trended to decline. Structural connectivity is widely weakened with age. Low-frequency NC-SC coupling in the PFC increased, while high-frequency coupling consistently decreased. Theta NC-SC couplings in the PFC were associated with memory changes. These results highlight the complex alterations in DMN information transmission across the lifespan, indicating the DMN’s potential as a neural marker for aging in the course of the adult lifespan.

It has been shown that low-frequency oscillations are linked to functional inhibition, and fast-band oscillations represent cortical activation^35^. Extending previous neuroimaging evidence in DMN, we detailed the multiple frequency bands neurophysiological changes across the adult lifespan. Specifically, our study revealed the gradual slowing of spontaneous activity frequency in the DMN during human adult lifespan. This change tendency was similar to previous research focusing on the whole-brain resting-state activities with EEG and MEG^35-37^. It has been shown that low-frequency oscillations are linked to functional inhibition, and fast-band oscillations represent cortical activation^38^. Moreover, the reduction of fast oscillations (alpha, beta, and gamma frequencies) and the general enhancement of slow rhythm (theta frequency) are the most common resting state EEG/MEG findings in patients with AD^39^. A previous research indicated that the age-related increase in theta power was associated with total-tau and phosphorylated tau levels in cerebrospinal fluid, suggesting that increased theta oscillations may serve as an early indicator of neurodegeneration^40^. Meanwhile, several neurotransmitter systems, including GABAergic, glutamatergic, and cholinergic neurons, contribute to the primary generation of high-frequency oscillations^41^. Overall, the change tendency in the DMN rhythms observed through MEG may reflect the brain aging processes throughout the lifespan. Simultaneously, structural connectivity within these subregions weakens with age, further revealing the detrimental impact of aging in the late adult lifespan on brain structure.

Our study has revealed significant insights into the coupling mechanisms within the DMN across adult lifespan and their associations with memory changes. Specifically, the observed increase in low-frequency NC-SC coupling (theta band) within the PFC is a biomarker in memory decline. More importantly, growing recent studies have demonstrated that interventions targeting high-frequency activity rather than low-frequency, such as game-frequency transcranial alternating current stimulation (tACS), can improve memory performance in aging adults, thereby underscoring the therapeutic potential of enhancing theta coupling^24^.

Our results also demonstrate that the coupling patterns within the PCC and temporal regions are distinct from those in the PFC. The consistent increase in PCC coupling across all frequency bands suggests a general enhancement of connectivity within this region, which may support its role in various cognitive functions beyond memory^43^. The increased interhemispheric connectivity in the temporal subregions further highlights the complex interplay between different brain regions in maintaining cognitive functions across adult lifespan^44^, consistent with previous findings that emphasize the importance of interhemispheric communication in cognitive resilience^45^. These findings illuminate the complex, frequency-dependent changes within the DMN across the adult lifespan, highlighting the importance of theta coupling in mitigating memory loss and suggesting potential interventions to enhance cognitive resilience in the aging population.

These findings illuminate the complex, frequency-dependent changes within the DMN across the adult lifespan, highlighting the importance of theta coupling in indicating memory loss and suggesting potential interventions to enhance cognitive resilience in the aging population.

### Limitation

While our study provides valuable insights into connectome alterations within the default mode network (DMN), several limitations should be acknowledged. First, by focusing primarily on white matter networks, our study offers an incomplete perspective on the effects of aging on brain structure, as changes in gray matter and microstructural integrity also play crucial roles in aging and cognitive decline. Additionally, our investigation is centered on memory-related alterations within the DMN, potentially overlooking its involvement in other cognitive domains, such as attention and executive function. A more comprehensive exploration of these mental functions is necessary to fully elucidate the role of the DMN in the aging process. Moreover, the cross-sectional nature of our study limits our ability to draw causal inferences between DMN alterations and cognitive decline. Longitudinal studies are essential to understand how these neural changes unfold over time and their direct impact on cognition. Finally, despite the breadth of our dataset, it may lack sufficient diversity in terms of genetic background, lifestyle factors, and comorbid conditions, potentially limiting the generalizability of our findings. Future research should aim to include more diverse samples and account for potential confounding variables such as mental health status and medication use to enhance the robustness and applicability of the results.

## Conclusion

Our study systematically elucidates the alterations in information transmission patterns within the DMN during aging, integrating high temporal resolution MEG, high spatial resolution functional MRI, and diffusion tensor imaging data. We discovered that aging enhances low-frequency (theta band) connectivity within DMN subregions, specifically between the PFC, temporal regions, and PCC while decreasing high-frequency (alpha, beta, gamma bands) connectivity. Structural connectivity within these subregions weakened with age, highlighting the detrimental impact of aging on brain integrity. Moreover, we found low-frequency NC-SC coupling (theta band) within the PFC increased with aging. In contrast, high-frequency coupling decreased, and PCC coupling values consistently increased across all frequencies. Notably, theta NC-SC in the PFC was associated with memory changes. These findings suggest that the DMN could serve as a potential neural marker for aging, providing valuable insights into the neural diversity underlying cognitive decline in older adults.

## DATA AVAILABILITY

MEG, fMRI, DWI data could be available on https://cam-can.mrc-cbu.cam.ac.uk/dataset/.

## CODE AVAILABILITY

Code will be available upon reasonable request.

## ACKNOWLEDGMENTS

Xiaobo Liu is supported by the China Scholarship Council. This research was supported by the National Programs for Brain Science and Brain-like Intelligence Technology of China (STI2030-Major Projects, No. 2021ZD0204004) and the National Natural Science Foundation of China (No. 82101559).

